# Single domain shark VNAR antibodies neutralize SARS-CoV-2 infection in vitro

**DOI:** 10.1101/2021.06.08.447530

**Authors:** Aziz Gauhar, Cyril V Privezentzev, Mykhaylo Demydchuk, Tanja Gerlza, Julia Rieger, Andreas J Kungl, Frank S Walsh, J Lynn Rutkowski, Pawel Stocki

## Abstract

Single domain shark VNAR antibodies can offer a viable alternative to conventional Ig-based monoclonal antibodies in treating COVID-19 disease during the current pandemic. Here we report the identification of neutralizing single domain VNAR antibodies selected against the SARS-CoV-2 spike protein derived from the Wuhan variant using phage display. We identified 56 unique binding clones that exhibited high affinity and specificity to the spike protein. Of those, 10 showed an ability to block both the spike protein receptor binding domain from the Wuhan variant and the N501Y mutant from interacting with recombinant ACE2 receptor *in vitro*. In addition, 3 antibody clones retained *in vitro* blocking activity when the E484K spike protein mutant was used. The inhibitory property of the VNAR antibodies was further confirmed for all 10 antibody clones using ACE2 expressing cells with spike protein from the Wuhan variant. The viral neutralizing potential of the VNAR clones was also confirmed for the 10 antibodies tested using live Wuhan variant virus in *in vitro* cell infectivity assays. Single domain VNAR antibodies due to their low complexity, small size, unique epitope recognition and formatting flexibility should be a useful adjunct to existing antibody approaches to treat COVID-19.

## Introduction

COVID-19 disease is caused by the transmission of severe acute respiratory syndrome coronavirus 2 (SARS-CoV-2) and is associated with global infections leading to a high mortality. The pandemic is predicted to circulate worldwide for the foreseeable future. SARS-CoV-2 is a member of coronavirus family that shares nearly 80% sequence homology with SARS-CoV (1, 2). Similar to SARS-CoV, the spike (S) protein expressed on the surface of the SARS-CoV-2 virion engages with host cells through its receptor angiotensin converting enzyme 2 (ACE2), which results in fusion of the viral envelope with the cell membrane and entry into host cells (3, 4). The extracellular domain of the S protein consists of the S1 and S2 subunits, which mediate receptor binding and membrane fusion, respectively. S1 harbors the receptor binding domain (S1-RBD), which directly contacts ACE2 (5), and the heavily glycosylated N-terminal domain (S1-NTD), which interacts with C-type lectin receptors (6, 7). Cell entry is driven by S2 and requires priming by cellular proteases that cleave the S protein between S1 and S2 subunits, and at S2’ site to allow membrane fusion (8, 9).

Intensive efforts to develop vaccines against SARS-CoV-2 have largely been successful, using a wide array of innovative and traditional approaches, including adenovirus vector- and mRNA-based vaccines (10). While the rollout of vaccination programs shows very high level of protection in immunized populations, establishing long-term safety and efficacy of vaccines will require continuous monitoring of protection especially against newly emerging variants. As vaccine-induced immunity can be less effective in older populations and immunocompromised individuals, and can potentially trigger a harmful response in some cases (11), then passive immunization via the delivery of purified antibodies could provide a solution (12). The most clinically advanced monoclonal antibodies target the S1-RBD and were either developed from convalescent patients infected with SARS-CoV-2 (13–15) or by immunizing humanized mice against the S protein (16, 17). Besides monoclonal antibodies, single domain antibodies from human and camelid origin that target S1-RBD have also shown potent virus neutralization activity *in vitro* (18–20).

The emergence of new variants is driven by mutation prone viral replication and selective mechanisms involving the human host. Thus, the emerging novel variants will be naturally selected for their ability to infect/reinfect the human population and escape the immune response. Consequently, vaccination and passive immunization with Ig-based antibodies may add to the selection pressure and the emergence of new variants with escape potential leading to decreased protection (21).

An alternative therapeutic approach using single domain shark VNAR antibodies could be of immense value. Due to their unique structure and characteristic protruding paratopes, VNAR antibodies can engage with buried cryptic epitopes that are not available to conventional Ig-based antibodies (22–24). VNARs are also not part of human immune response, thus would be more likely to retain neutralizing activity against new variants emerging naturally under selection pressure from Ig-based antibodies in an immunized or previously infected population. In addition, VNAR antibodies possess advantageous properties including small size, less complex structure, and a desirable chemical and thermal stability profile that allows for ease of manufacturing, storage without low temperature refrigeration, and a variety of formulation options including the inhalation route.

Here, we report the identification of neutralizing single domain VNAR antibodies derived by bio-panning against the Spike protein from the SARS-CoV-2 Wuhan variant. The screens identified 10 VNAR antibodies that were shown in biochemical and cells-based assays to be highly effective in blocking the interaction of recombinant Spike protein with ACE2 receptor and were further confirmed to block live Wuhan variant virus using *in vitro* assays. All 10 clones retained high affinity binding and blocking activity against the S1-RBD N501Y mutant, and 3 showed neutralizing activity against the S1-RBD E484K mutant. The VNAR antibodies directed against the SARS-CoV-2 spike protein expand the molecular toolbox of novel therapeutic approaches against COVID-19.

## Methods and Materials

### Protein expression and purification

The S1 subunit (residues 16-685) and S1-RBD domain (residues 319-541) of SARS-CoV-2 spike protein (NCBI Accession: YP 009724390.1) were synthesized and cloned into the pFUSE expression vector (InvivoGen). Both constructs contained an AVI-tag and a 6xHis tag at the C-terminus separated by a single G_4_S linker. The ACE2 (UniProt entry: Q9BYF1) ectodomain (18-740) was synthesized with FLAG and 6xHis tags at the C-terminal end separated by a single G_4_S linker before cloning into the pFUSE vector. S1, S1-RBD and ACE2 recombinant proteins were expressed in Expi293 cells for 5 days following transient transfection using Expifectamine 293 Transfection Kit (Thermo Fisher Scientific). Cell cultures were centrifuged at 4500*g* for 40 min, supernatants were collected and filtered on 0.45 μm membranes before being loaded on a HisTrap Excel column (Cytiva). After extensive washing, the proteins were eluted with 500 mM imidazole, concentrated and buffer exchanged to PBS pH 7.4 using HiPrep 26/10 desalting columns. The quality of the purified proteins was assessed by sodium dodecyl sulfate polyacrylamide gel electrophoresis (SDS-PAGE) and analytical size exclusion chromatography (SEC) using a Superdex 200 Increase 10/300 GL analytical column (Cytiva).

### Phage display selection campaign

Two different VNAR libraries based on type II VNAR nurse shark single domain antibody sequences were used for selections. The OSX3 library contained variable CDR3 lengths from 11 to 18 amino acids with a flexible position of the canonical cysteine in the CDR3 loop (25). The OSX6 library contained a single length CDR3 of 14 amino acid residues with a centrally fixed canonical cysteine within approximately 10^3^ framework variants identified from naïve and immune libraries by next-generation sequencing. Phage selections were performed against the S1-RBD domain (Trenzyme) and S1 subunit of SARS-CoV-2 spike protein, which were biotinylated using EZ-Link Sulfo-NHS-Biotin reagent (Pierce). PEG/NaCl precipitated phage libraries blocked in 5% BSA in PBS were deselected with streptavidin magnetic beads prior to bio-panning against biotinylated S1 or S1-RBD. Phages bound to biotinylated target protein were captured by streptavidin coupled Dynabeads (Thermo Fisher Scientific), washed and eluted in 100 nM triethylamine. Eluted phages were adjusted to neutral pH and propagated in TG1 *E. coli*. M13KO7 helper phage was used to induce phage production for subsequent rounds of selection. The amount of antigen used in each subsequent selection round was 100, 50, and 25 nM for round 1, 2 and 3, respectively.

### Phage ELISA

Individual clones were picked from agar plates and grown at 37°C with shaking in a 96-well block in 2YT media supplemented with 2% glucose and 100 μg/ml ampicillin until visible growth occurred. The cultures were super-infected with M13K07 helper phage (New England Biolab) for 1 hr and the media was replaced with 2YT media supplemented with 100 μg/ml ampicillin and 50 μg/ml kanamycin. After 16 hr at 30°C, supernatants were collected by centrifugation and blocked in a final concentration of 2.5% milk in PBS with 0.1% Tween-20 (PBST) for 1 hr at room temperature (RT). High binding 96 well microplates (Greiner) were coated with 100 μl (5 μg/ml) of purified S1 or S1-RBD or commercial human serum albumin (HSA; Sigma-Aldrich) and incubated overnight at 4°C. Plates were blocked with 2.5% milk in PBST for 1 hr at RT. Blocked phage supernatants were transferred to blocked microplates and incubated for 1 hr followed by washing with PBST on a plate washer (BioTek). Anti-M13-HRP antibody (Sino Biological) was used for detection at 1:4000 dilution in blocking buffer. The reaction was developed with 1-Step Ultra TMB-ELISA Substrate Solution (Thermo Fisher Scientific), stopped with 1% HCl and absorbance was measured at 450 nm on VarioSkan plate reader (Thermo Fisher Scientific).

### VNAR-Fc formatting

Unique VNAR clones that showed specific binding to the antigen in phage ELISA were identified by Sanger sequencing and cloned into the pFUSE expression vector. Expi293 cells were transiently transfected as VNAR human Fc fusions (VNAR-hFc) and produced following the manufacturer’s manual. After 5 days growth, cell cultures were centrifuged at 2,000 rpm for 10 min, supernatants were filtered using 0.22 μm membrane filters and loaded onto HiTrap MabSelect SuRe columns (Cytiva) pre-equilibrated against PBS, pH 7.4. Protein A bound antibodies were eluted with 0.1 M Glycine, pH 3.5 and the buffer exchanged to PBS, pH 7.4 using HiPrep 26/10 desalting column (Cytiva). Purity of the purified protein samples was determined by analytical SEC and SDS-PAGE.

### Binding ELISA

To determine the binding EC_50_ of VNAR clones, high binding 96 well microplates (Greiner) were coated overnight at 4°C with 100 μl of S1, S1-RBD or HSA at 5 μg/ml and S1-RBD E484K and S1-RBD N501Y at 1 μg/ml. Plates were blocked for 1 hr at RT with 5% bovine serum albumin (BSA) in PBS before a 1 hr incubation with serially diluted VNAR-hFc antibodies or ACE2. The plates were washed, and binding was detected with an anti-human IgG (Fc specific) (1:5000 dilution, Sigma-Aldrich, A0170) or anti-FLAG (1:1000 dilution, Sigma-Aldrich, A8592) HRP-conjugated antibodies for VNAR-hFc antibodies or ACE2, respectively. The signal was developed with 1-Step™ Ultra TMB-ELISA substrate solution (Thermo Fisher Scientific) and the reaction was stopped with 1% HCl. Absorbance measured at 450 nm and EC_50_ values were calculated by 4-parametric non-linear regression using GraphPad Prism 8.0.

### Blocking ELISA

The IC_50_ was determined by ELISA as above except that serial dilutions of the VNAR-hFc antibodies were premixed with 1.4 nM of purified ACE2 prior to incubation in coated microtiter plates. ACE2 binding was measured with an anti-FLAG HRP conjugated antibody (Sigma-Aldrich, A8592) diluted 1:1000.

### Epitope binning

Pairwise competition ELISA was used to perform epitope binning. Individual VNAR-hFcs were used as primary antibodies at 50 nM concentration and 100 μl volume to coat high binding 96 well microplates overnight at 4°C. After blocking in 5% BSA in PBS for 1 hr, 50 nM biotinylated S1-RBD alone or premixed with 500 nM secondary (competitor) VNAR-hFc antibodies were added to the plate pairwise. Binding of biotinylated S1-RBD to coated VNAR-hFc was detected by streptavidin-HRP (Merck Millipore, 18-152). The signal intensities were collected for each primary VNAR-hFc antibody without competitor and compared to the signal when a competitor antibody was present. If the signal remained the same, it was defined as non-competitive, whereas if the signal was decreased by at least 20% it was defined as competitive.

### Flow cytometry

Expi293 cells were transiently transfected with ACE2 cloned into the pCMV3-C-GFPSpark expression vector (Sino Biological). Cells were incubated at 37°C in 8% CO_2_ shaking at 350 RPM for 48 hr. The collected cells were blocked in 2% BSA in PBS for 30 min at 4°C and then transferred to a 96 well V-shape microplate (Greiner) at a density of 100,000 per well. S1 protein was biotinylated at the AVI-tag using the BirA biotin-protein ligase kit (Avidity Biosciences) and premixed at 5 nM with serial dilutions of VNAR-hFc antibodies for 1 hr at 4°C. Binding of S1 to ACE2 transfected cells was measured with an Alexa Fluor-647-conjugated streptavidin (Invitrogen, S21374) diluted 1:500 using a CytoFLEX flow cytometer (Beckman Coulter). After gating on cells expressing ACE2-GFP fusion protein in FL1 channel, median fluorescence intensity (MFI) of cells positive for S1 binding was used to determine IC_50_ by 4-parametric non-linear regression analysis using GraphPad Prism 8.0.

### Viral neutralization

All studies with live virus were conducted in a certified BSL3 laboratory. Vero CCL81 cells were seeded at a cell density of 100,000 cells per well in 48-well plates and incubated at 37°C in serum free OptiPro SFM (Thermo Fisher Scientific) for 24 hr before infection. The SARS-CoV-2 Wuhan strain propagated in Vero CCL81 cells was preincubated with or without VNAR-hFc antibodies for 10 min in OptiPro medium before incubation of prewashed Vero CCL81 cells with and without the mix for 1 hr at 37°C. Supernatants were transferred to Eppendorf tubes and inactivated with 560 μl of AVL buffer from QIAamp Viral RNA Mini Kit (Qiagen). The cells were washed and further incubated at 37°C for 48 hr before supernatant was again harvested and the virus inactivated in AVL buffer. Cell death was assessed by phase microscopy at the end of the experiment.

Inactivated supernatants were transferred from the BSL3 lab and mRNA was isolated using the QIAamp Kit according to manufacturer’s protocol. Briefly, 560 μl absolute ethanol was added and loaded onto columns. After washes with AW1 and AW2 buffers, RNA was collected using 40 μl of Ambion nuclease free water (Thermo Fisher Scientific). Total RNA (5 μl) was used for cDNA synthesis and qPCR was performed in one step using QuantiTect Probe RT-PCR (Qiagen) on a StepOnePlus System (Applied Biosystems). The qPCR primers (Eurofins) were as follows: N1 forward GAC CCC AAA ATC AGC GAA AT, N1 reverse TCT GGT TAC TGC CAG TTG AAT CTG and N1 Probe FAM-ACC CCG CAT TAC GTT TGG TGG ACC-BHQ1. To assess RNA quality RNase P Primers were used: RP Forward AGA TTT GGA CCT GCG AGC G, RP Reverse GAG CGG CTG TCT CCA CAA GT and RP Probe FAM – TTC TGA CCT GAA GGC TCT GCG CG – BHQ-1. qPCR primers were used at 0.4 μM with the probe at 0.2 μM and samples were incubated at 50°C for 30 min, heated to 95°C for 15 min, followed by 45 cycles of 95°C for 3 sec and 55°C for 30 sec. C_t_ values obtained after 48 hr were subtracted from C_t_ values at the time of infection and the data normalized between untreated cells exposed to the virus set to 100% and cells that were not exposed to the virus (sham) set to 0%.

## Results

### Isolation of single domain VNAR antibodies to SARS-CoV-2 spike protein

Three consecutive rounds of in-solution bio-panning against biotinylated S1 and S1-RBD of SARS-CoV-2 spike protein were performed using two VNAR semi-synthetic phage libraries, OSX3 and OSX6. A strong enrichment in phage titer was observed from selection round 1 to round 3 (Fig. 1A). The proportion of clones binding to S1 and S1-RBD increased substantially from round 2 to round 3 as tested by phage ELISA (Fig. 1B). To assess the diversity and identify individual VNAR antibodies, the clones that showed specific binding to S1 or S1-RBD were sequenced. The diversity of clones was higher in round 2 in comparison to round 3 for both libraries (Fig. 1C). From round 3 outputs, 35 and 20 unique VNARs were identified for S1-RBD and S1 selections, respectively. Whereas from round 2 outputs, 59 and 35 unique VNARs were confirmed for S1-RBD and S1 selections, respectively.

**Figure 1.**
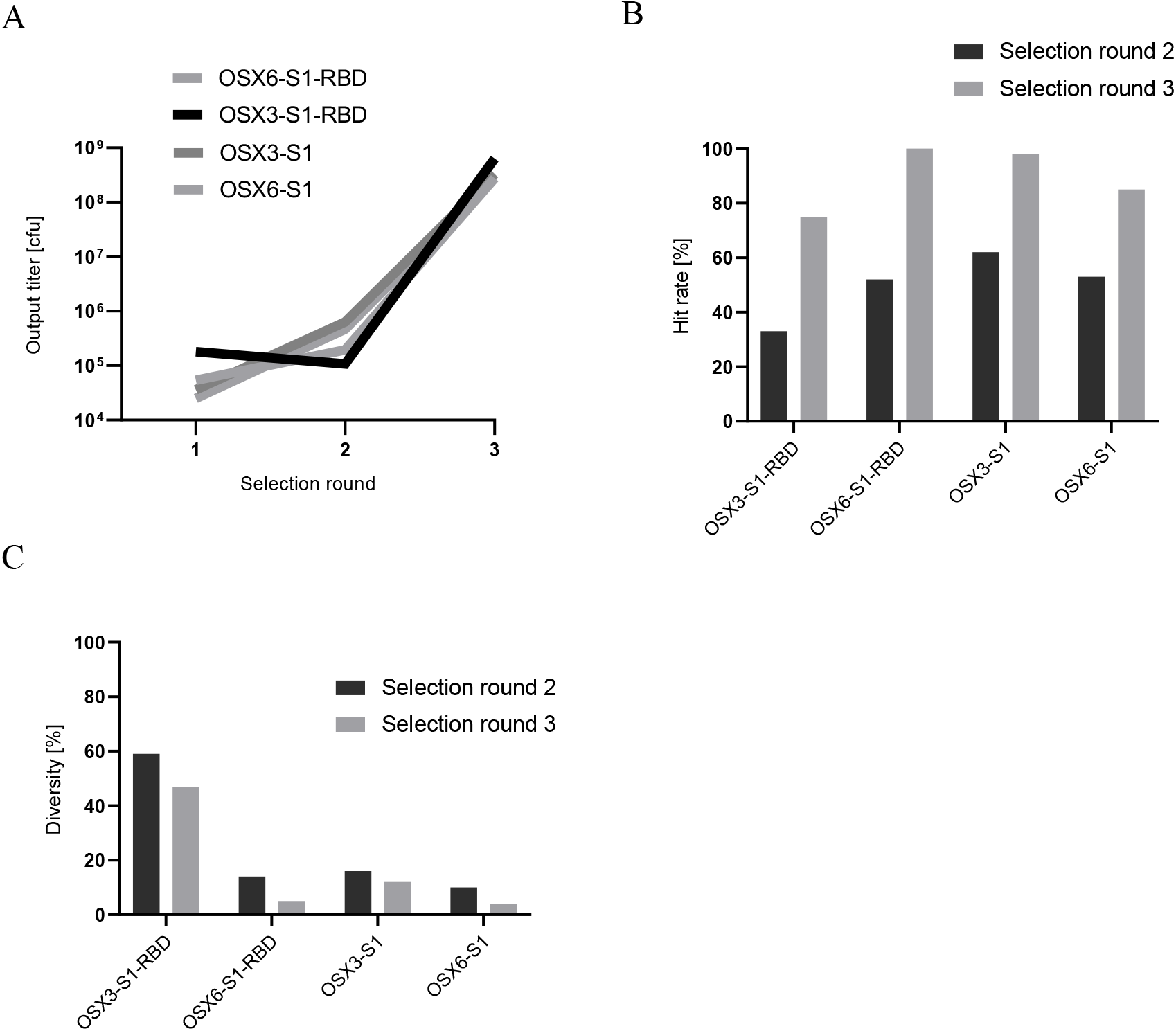
Isolation of SARS-CoV-2 specific VNARs. **(A)** Output titers for OSX3 and OSX6 libraries were calculated individually for each round of selection and presented as colony forming units (cfu) for the selection campaigns with S1 or S1-RBD. **(B)** Hit rate for rounds 2 and 3 determined by phage ELISA and expressed as the percentage of VNAR clones that specifically bound to S1-RBD or S1 with at least 3-fold signal over negative control. **(C)** VNARs that bound to S1 were sequenced and diversity was calculated as the percentage of unique clones after each round of selection.

In total 149 unique VNARs were reformatted to bivalent antibodies by fusing to human IgG1 Fc. The VNAR human Fc fusions (VNAR-hFc) were subsequently expressed in Expi293 cells and crude culture supernatants were used in binding ELISA for S1-RBD and S1 (Fig. 2 - shows representative 10 clones that were revealed to possess neutralizing activity in follow-up experiments). Of the reformatted VNAR-hFc antibodies, 60 retained strong binding to S1-RBD and/or S1 with OD >0.3 (approximately 3-4-fold over negative control) and 4 clones that showed nonspecific binding to HSA were removed from further analysis. Out of 56 specific clones, 37 and 19 originated from OSX3 and OSX6 library, respectively. Nine of the 56 clones showed binding to S1 but not S1-RBD suggesting that the epitope was present within S1-NTD fragment.

**Figure 2.**
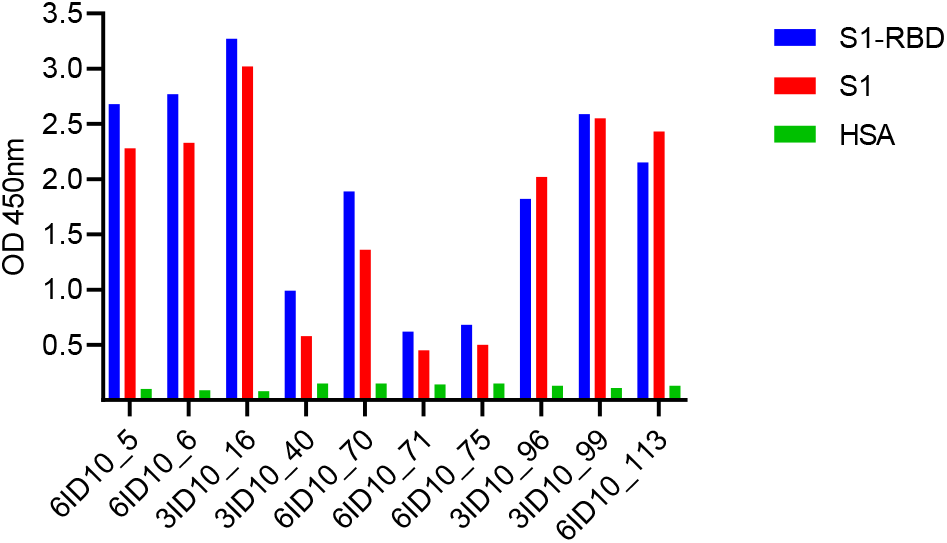
Binding ELISA of VNARs after reformatting to bivalent VNAR-hFc antibodies. VNAR-hFc antibodies were expressed in small scale and crude supernatant samples were tested by ELISA for binding to S1-RBD, S1 and HSA. Data presented as OD measured at 450nm.

All specific VNAR-hFc antibodies were expressed at medium scale and purified by protein A chromatography. Final antibody purity was >90% as determined by SEC and SDS-PAGE. Binding affinity of the purified VNAR-hFc antibodies to S1-RBD and S1 was assessed by ELISA. Of the 56 clones tested, 47 showed high EC_50_ binding affinities (<10nM) including the selected 10 clones with neutralizing activity (Fig. 3A and B, and Table 1). Binding of these 10 VNAR-hFc antibodies was assessed against two key mutations in S1-RBD region, N501Y and E484K found in the newly emerged SARS-CoV-2 variants (26, 27). Binding of VNAR-hFc antibodies to S1-RBD N501Y remained comparable to S1-RBD from the original Wuhan variant except for 3ID10_16, which increased EC_50_ affinity by approximately 90-fold (Fig. 3B and C, and Table 1), whereas binding to S1-RBD E484K was reduced for all of the tested clones in comparison to S1-RBD (Fig. 3D and Table 1).

**Figure 3.**
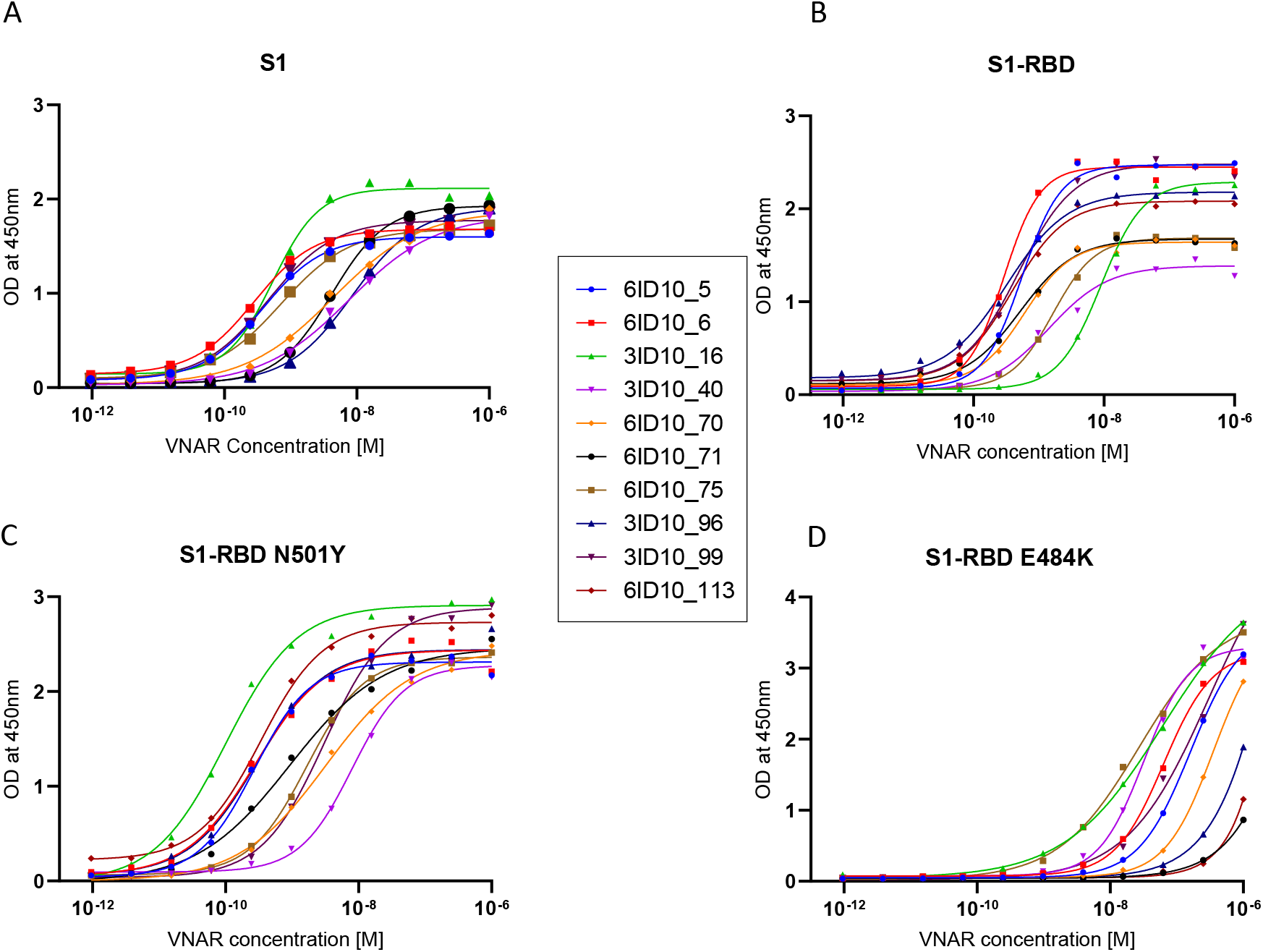
ELISA EC_50_ binding affinity of select VNAR-hFc antibodies to Wuhan and mutant spike proteins. ELISA was used to determine EC_50_ values for individual VNAR-hFc antibodies to **(A)** S1, **(B)** S1-RBD, **(C)** S1-RBD N501Y, and **(D)** S1-RBD E484K. Microplates were coated with different spike proteins and incubated with serially diluted VNAR-hFc antibodies followed by detection with HRP conjugated anti-human Fc antibody. The developed plates were read at 450nm and OD was used for 4-parametric non-linear regression model to calculate EC_50_ values (see Table 1).

**Table 1:** ELISA-based EC_50_ binding affinity shown for select VNAR-hFc antibodies and ACE2 against S1, S1-RBD, S1-RBD N501Y and S1-RBD E484K. The values were calculated using 4-parametric non-linear regression model and presented as molar values. n.a. – data not available

| VNAR-hFc | ELISA EC<sub>50</sub> [M] |  |  |  |
| --- | --- | --- | --- | --- |
|  | S1 | S1-RBD | S1-RBD N501Y | S1-RBD E484K |
| 6ID10_5 | 3.8E-10 | 5.5E-10 | 2.7E-10 | 1.6E-07 |
| 6ID10_6 | 2.8E-10 | 2.9E-10 | 2.9E-10 | 6.4E-08 |
| 3ID10_16 | 5.4E-10 | 8.9E-09 | 1.0E-10 | 6.3E-08 |
| 3ID10_40 | n.a. | 1.4E-09 | 8.0E-09 | 3.2E-08 |
| 6ID10_70 | 7.6E-09 | 6.0E-10 | 3.2E-09 | >1E-07 |
| 6ID10_71 | 4.4E-09 | 5.2E-10 | 9.3E-10 | >1E-06 |
| 6ID10_75 | 4.1E-09 | 1.7E-09 | 1.7E-09 | 2.7E-08 |
| 3ID10_96 | 7.2E-10 | 3.2E-10 | 3.0E-10 | >1E-06 |
| 3ID10_99 | 8.5E-09 | 4.9E-10 | 3.1E-09 | 3.1E-07 |
| 6ID10_113 | 4.6E-10 | 3.9E-10 | 3.4E-10 | >1E-06 |
| ACE2 | n.a. | 2.8E-09 | 1.5E-09 | 2.5E-09 |

To assess the epitope coverage of the spike protein by the selected 10 VNAR antibodies, epitope binning ELISA was performed. ELISA plate was coated with the primary VNAR-hFc antibody at 50 nM concentration, followed by incubation with either 50 nM biotinylated S1-RBD alone or a mixture of 50 nM biotinylated S1-RBD and 500 nM secondary VNAR-hFc antibody. After incubation, the plate was washed and streptavidin-HRP was used for detection of S1-RBD binding to immobilized primary antibody. Each primary antibody was tested against each secondary antibody for competition to S1-RBD binding and signal for no-secondary VNAR-hFc antibody was used as a reference control. Competition was defined as a signal decrease of at least 20% compared to the reference and indicated an overlap in the epitope bin between the antibodies, whereas the lack of competition suggested a unique bin. Overall, two bins were identified, with 9 out of 10 antibodies showing a shared overlapping bin and 1 antibody 3ID10_99 had a unique bin (Fig. 4).

**Figure 4.**
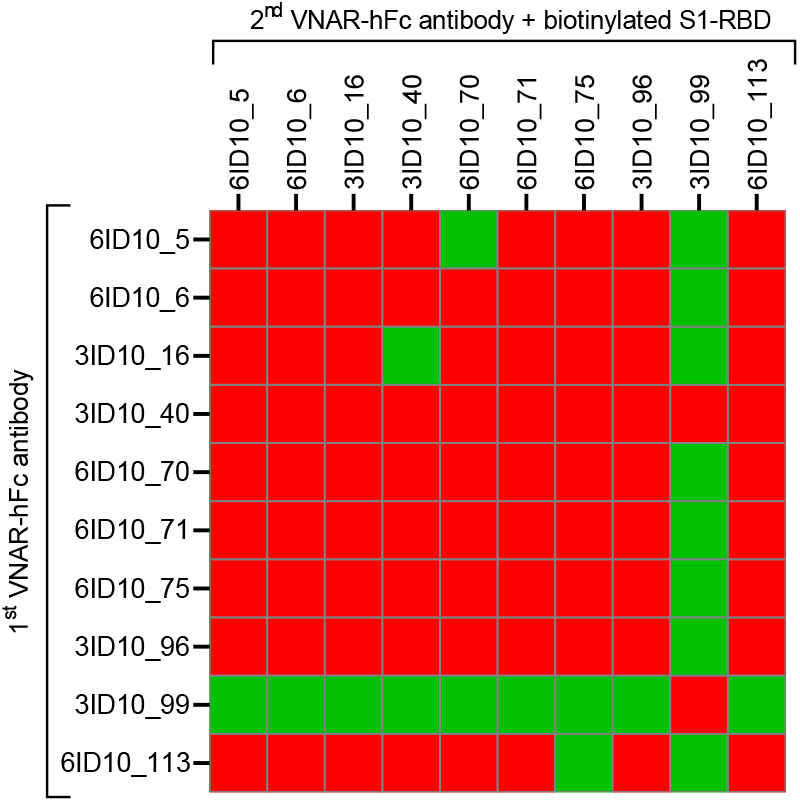
Epitope binning of VNAR-hFc antibodies by competition ELISA. Primary VNAR-hFc antibodies were used to coat ELISA microplates, followed by addition of S1-RBD or premixed biotinylated S1-RBD and secondary VNAR-hFc antibodies as potential competitor. Binding of biotinylated S1-RBD to immobilized VNAR-hFc antibodies was detected by streptavidin-HRP. Signal for wells lacking secondary VNAR-hFc antibody was used as a reference control for each primary antibody. Signal decrease of at least 20% in comparison to reference control indicated a competition marked in red and suggested overlapping epitope. Green color shows the lack of competition, thus a unique epitope.

### Inhibition of spike protein interaction with ACE2

The binding EC_50_ affinities of S1-RBD Wuhan variant and the N501Y and E484K mutants to ACE2 were measured to set up a comparative *in vitro* inhibitory assay. Only minor differences in ELISA EC_50_ affinities were observed for binding to ACE2 receptor between S1-RBD and the mutants (Fig. 5 and Table 1). S1-RBD N501Y showed higher EC_50_ affinity by approximately 2-fold whereas S1-RBD E484K was unchanged when compared to S1-RBD (Fig. 5 and Table 1). Purified VNAR-hFc antibodies were then tested in an inhibition ELISA for their ability to block the interaction of ACE2 with SARS-CoV-2 S1, S1-RBD or the mutants. The plates were coated with S1, S1-RBD or the mutants and incubated with serial dilutions of VNAR-hFc antibodies premixed with ACE2 at 1.4 nM. Out of 56 VNAR-hFc antibodies tested, 10 efficiently blocked the binding of ACE2 to S1 and S1-RBD with calculated IC_50_ values ranging from 2.1 to 36.5 and 7.5 to 92 nM, respectively (Fig. 6A and B, and Table 2). All 10 clones retained blocking ability for S1-RBD N501Y and 8 of these showed similar blocking IC_50_ values when compared to S1-RBD. Clone 3ID10_40 showed reduced and clone 3ID10_96 showed improved IC_50_ inhibitory potential by approximately 22 and 20-fold, respectively (Fig. 6B and C, and Table 2). Two antibodies 3ID10_16 and 6ID10_75 blocked S1-RBD E484K with nearly identical IC_50_ when compared to S1-RBD (Fig. 6D and Table 2). Antibody clone 3ID10_99 showed inhibitory activity in the ELISA assay; however, the curve did not fit non-linear regression analysis and did not produce a precise IC_50_ value, which was estimated to be 1-10 nM.

**Figure 5.**
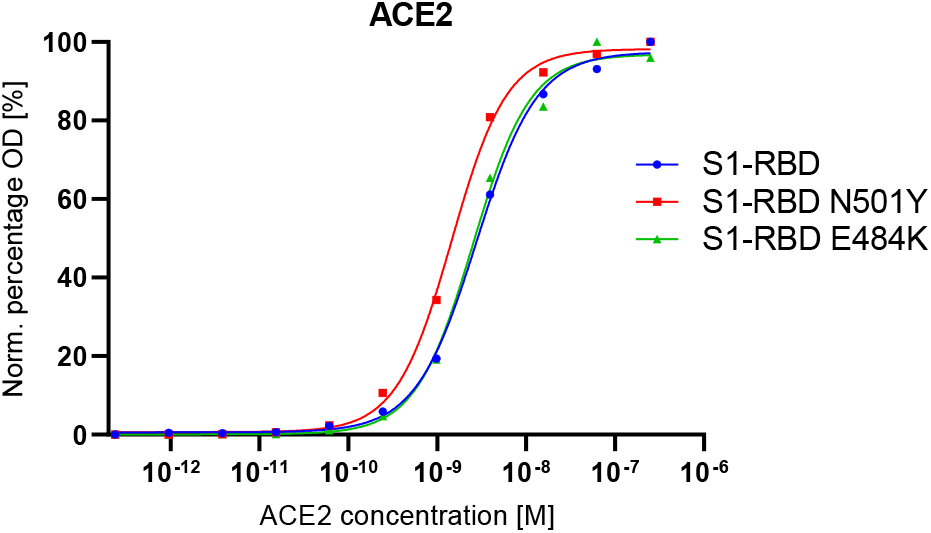
ELISA-based EC_50_ binding affinity of spike mutants to ACE2. ELISA was used to determine EC_50_ values for ACE2 binding to S1-RBD, S1-RBD N501Y or S1-RBD E484K. Microplates were coated with spike proteins and incubated with serially diluted ACE2 followed by detection with anti-FLAG-HRP used for ACE2 detection. The developed plates were read at 450nm and normalized OD was used for 4-parametric non-linear regression model to calculate EC50 values (see Table 1).

**Figure 6.**
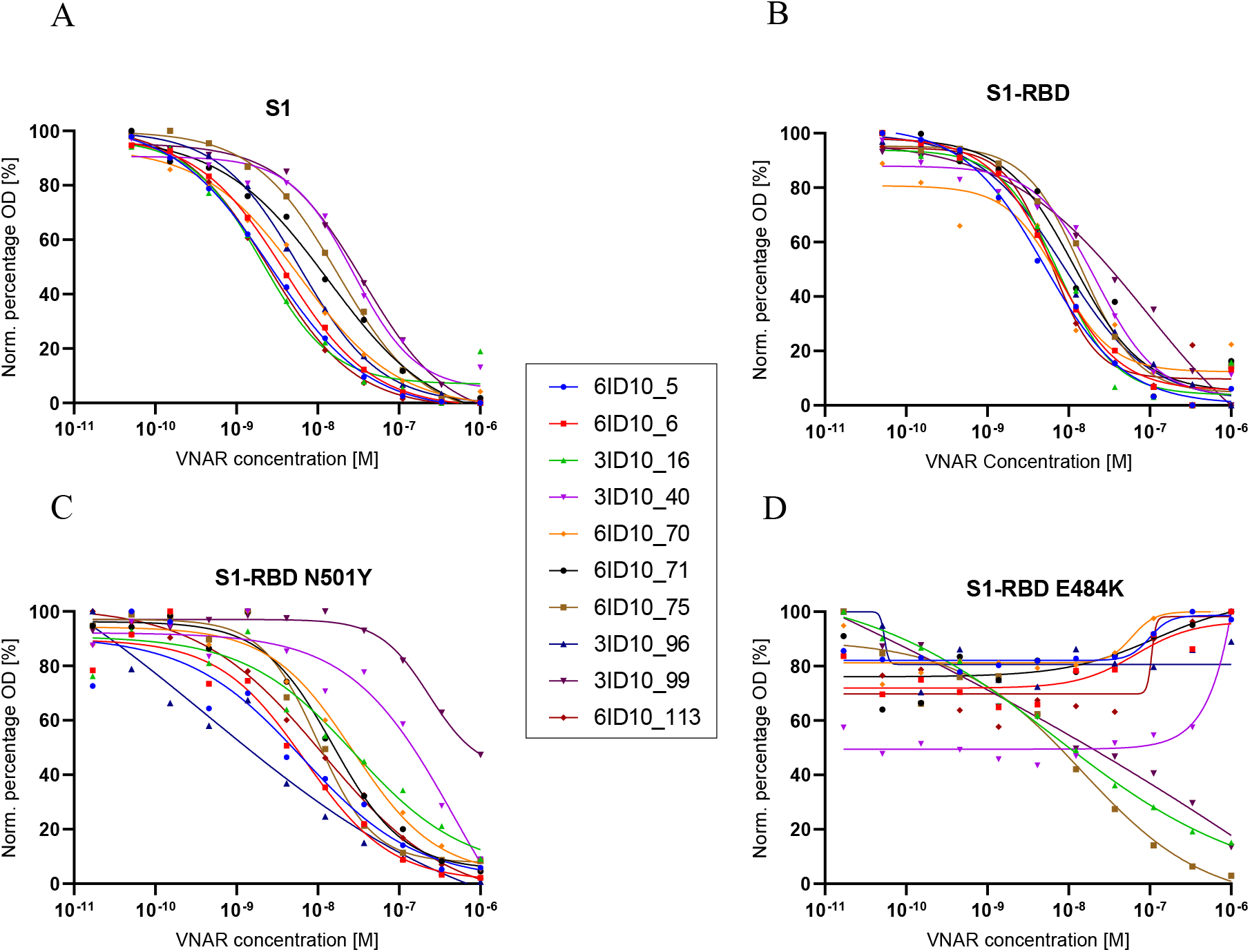
ELISA-based IC_50_ blocking activity of select VNAR-hFc antibodies. ELISA was used to determine IC_50_ blocking values for individual VNAR-hFc antibodies against **(A)** S1, **(B)** S1-RBD, **(C)** S1-RBD N501Y, and **(D)** S1-RBD E484K. Microplates were coated with spike proteins and incubated with serially diluted VNAR-hFc antibodies premixed with ACE2 at 1.4 nM concentration followed by detection with anti-FLAG-HRP used for ACE2 detection. The developed plates were read at 450nm and normalized OD was used for 4-parametric non-linear regression model to calculate IC_50_ values (see Table 2).

**Table 2:** ELISA-based IC_50_ blocking activity shown for select VNAR-hFc antibodies against S1, S1-RBD, S1-RBD N501Y and S1-RBD E484K. Cell-based blocking activity was performed with transiently transfected cells expressing ACE2 and VNAR-hFc antibodies against S1 or S1-RBD and was assessed by flow cytometry. The values were calculated using 4-parametric non-linear regression model and presented as molar values. n.d. - not detected, * – approximated value due to poor fit to non-linear regression model.

| VNAR-hFc | ELISA IC<sub>50</sub> [M] |  |  |  | Cell assay IC<sub>50</sub> [M] |  |
| --- | --- | --- | --- | --- | --- | --- |
|  | S1 | S1-RBD | S1-RBD N501Y | S1-RBD E484K | S1 | S1-RBD |
| 6ID10_5 | 2.6E-09 | 5.0E-09 | 6.5E-09 | n.d. | 5.9E-08 | 1.5E-07 |
| 6ID10_6 | 3.9E-09 | 6.4E-09 | 6.9E-09 | n.d. | 9.4E-08 | 1.2E-08 |
| 3ID10_16 | 2.1E-09 | 7.5E-09 | 2.8E-08 | 6.9E-09 | 9.6E-08 | 7.5E-08 |
| 3ID10_40 | 2.8E-08 | 2.2E-08 | 5.0E-07 | n.d. | 7.7E-07 | 4.6E-07 |
| 6ID10_70 | 6.1E-09 | 7.7E-09 | 2.7E-08 | n.d. | 2.1E-07 | 1.7E-07 |
| 6ID10_71 | 1.4E-08 | 1.2E-08 | 1.5E-08 | n.d. | 2.8E-07 | 9.9E-07 |
| 6ID10_75 | 1.8E-08 | 1.5E-08 | 9.9E-09 | 1.5E-08 | 2.1E-07 | 2.1E-06 |
| 3ID10_96 | 6.0E-09 | 8.8E-09 | 4.4E-10 | n.d. | 1.8E-07 | 6.1E-07 |
| 3ID10_99 | 3.7E-08 | 9.2E-08 | 2.3E-07 | 1-10E-08* | 8.3E-07 | 3.1E-07 |
| 6ID10_113 | 2.5E-09 | 6.1E-09 | 1.2E-08 | n.d. | 8.6E-08 | 7.9E-08 |

The 10 VNAR-hFc antibodies that blocked binding of ACE2 to S1 or S1-RBD in the competition ELISA were further tested in a cell-based assay. Expi293 cells were transiently transfected with ACE2 fused to GFP, which allowed precise gating of transfected cells during the flow cytometry analysis. GFP was fused to the C-terminal end of cytoplasmic domain of ACE2 to minimize potential consequences to the structure of the ectodomain, thus the ability to interact with recombinant S1 or S1-RBD proteins. The transiently transfected cells were incubated with biotinylated S1 or S1-RBD proteins at 5 nM concentration in the presence of serially diluted VNAR-hFc antibodies. Site-specific biotinylation of C-terminal AVI-tag present in the S1 and S1-RBD constructs minimized potential steric hindrance upon binding from the fluorophore-conjugated streptavidin used for detection. The results showed that the tested VNAR-hFc antibodies were able to block S1 and S1-RBD binding to the cells expressing ACE2, which was in line with the *in vitro* competition by ELISA. The calculated IC_50_ values ranged from 40 nM to 1.6 μM, and from 75 nM to 1.8 μM for S1 and S1-RBD, respectively (Fig. 7 and Table 2).

**Figure 7.**
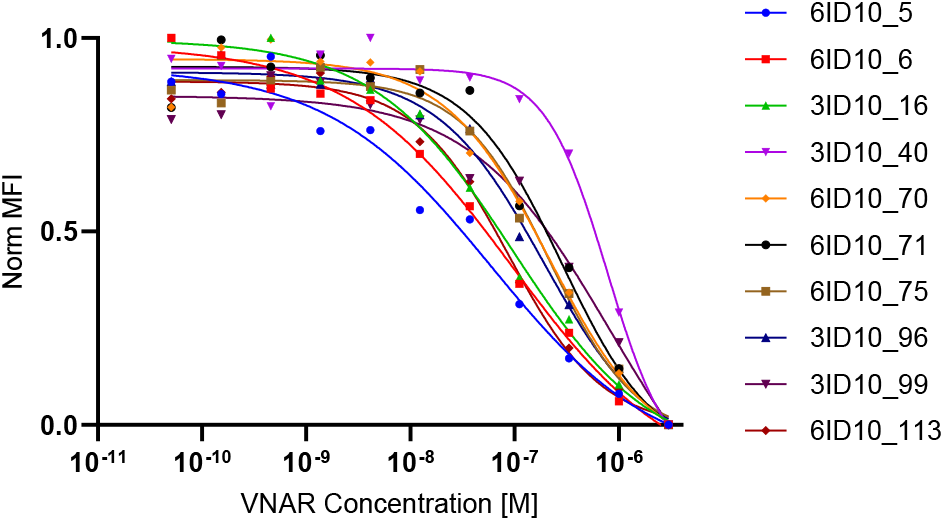
Cell-based IC_50_ blocking activity of select VNAR-hFc antibodies. Flow cytometry was used to determine IC_50_ blocking values for individual VNAR-hFc antibodies against S1. ACE2 with GFP fusion was transiently expressed in Expi293 cells for 48 hours following transfection. Cells were blocked before incubation with serially diluted VNAR-hFc antibodies premixed with biotinylated S1 at 5 nM concentration. Cells were washed and binding of S1 was detected with Alexa-Fluor-647-conjugated streptavidin. Cells expressing ACE2 were gated for GFP fluorescence in FL1 channel. Median fluorescence intensity (MFI) of transfected cell population was used to determine IC_50_ by 4-parametric non-linear regression model (see Table 2).

### Virus neutralization assay

The panel of 10 VNAR-hFc antibodies was further assessed for viral blocking using live SARS-CoV-2 Wuhan strain. The infection level of Vero CCL81 cells was determined by qPCR at the 48 hr timepoint post exposure. The assay was performed semi-quantitatively, using 5, 10, and 50 μg/ml (65, 130, and 650 nM) concentrations. A VNAR-hFc isotype control was included in the assay (used at 500 μg/ml (6.5 μM)) that showed no activity, whereas the 10 antibody clones tested were all able to prevent viral infection, of which 6 clones (6ID10_5, 6ID10_6, 6ID10_70, 6ID10_71, 6ID10_75, and 3ID10_99) showed ≥ 90% virus neutralization (Fig. 8). Consequently, it was concluded that applied screening and selection of clones that relied on *in vitro* and cell-based assay can accurately predict inhibitory potential when tested with the live virus.

**Figure 8.**
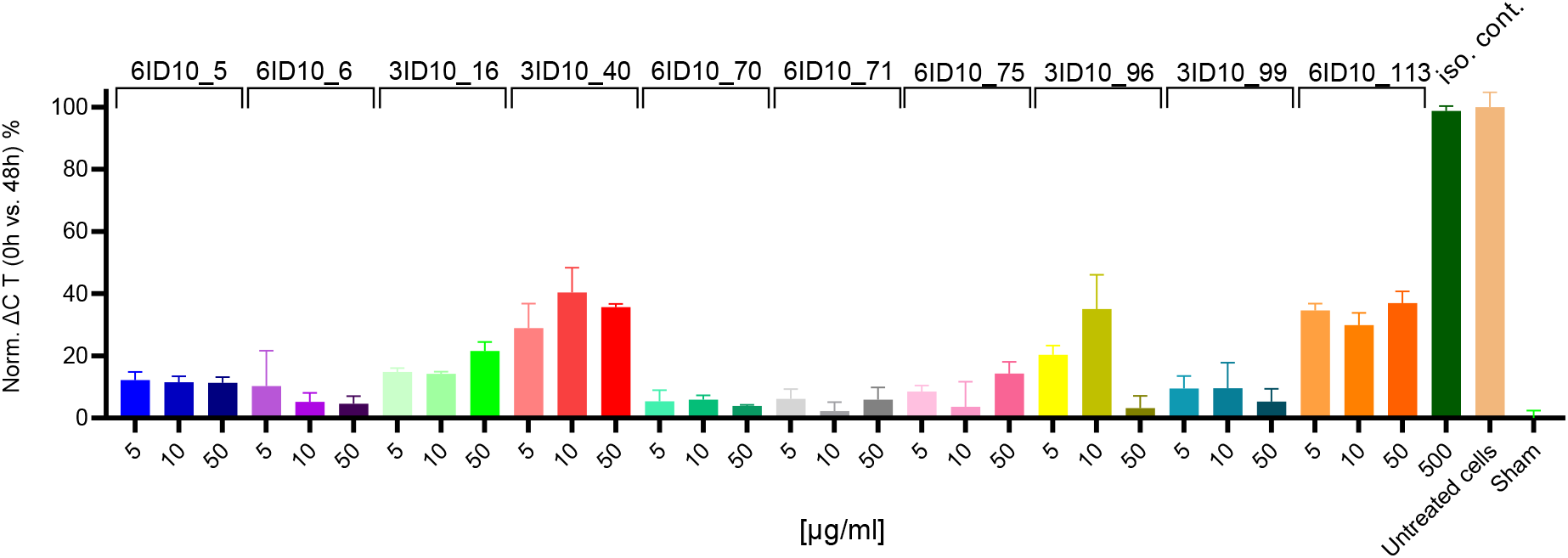
Live SARS-CoV-2 virus neutralization by VNAR-hFc antibodies. Live SARS-CoV-2 virus neutralization mediated by VNAR-hFc antibodies was performed using Vero CCL81 cells. SARS-CoV-2 Wuhan strain was preincubated with VNAR-hFc antibodies at 5, 10, and 50 μg/ml (65, 130, and 650 nM) or VNAR isotype control at 500 μg/ml (6.5 μM) concentration for 48 hr and viral load was determined using qPCR. Blocking activity was determined relative to cells treated with the virus in the absence of VNAR antibodies. Plotted ΔC_t_ values represent the mean of three technical repeats, ±SD.

## Discussion

In early 2020 the WHO officially declared COVID-19 as a pandemic. Since then, joint efforts by both public and private research institutions have led to the generation of an unprecedented amount of data regarding disease pathogenesis, epidemiology, virus structure and phylogenetics, its interaction with the host and most importantly the development of vaccines and other therapeutics such as antibodies. It has been found that the RBD fragment of the SARS-CoV-2 Spike protein binds with high affinity to the human and bat ACE2 receptors (28). As such the protein and specifically the S1-RBD domain became the focus of R&D efforts for developing vaccines and antibody therapeutics. A number of potent vaccines have been approved for human use by a large number of national drug regulatory authorities (29, 30). In addition, therapeutic antibodies isolated by a number of strategies have been approved and are currently used in the severely ill (16, 31, 32). Single domain antibody platforms of human and camelid (VHH) sources also showed promise in generating potent virus inhibitors (18, 19, 33, 34). In the current study, we have assessed the ability of the shark single domain VNAR platform to deliver target specific, neutralizing antibodies against SARS-CoV-2.

We targeted both the S1-RBD domain and the whole S1 subunit of the spike protein to isolate VNAR antibodies from our libraries using phage display. Both selection campaigns led to the generation of 56 target specific binders. Antibodies isolated from the S1-RBD selection campaign bound specifically to the S1-RBD domain, as did the majority of antibodies isolated using the S1 subunit. Only 9 antibody clones interacted with non-RBD regions of the S1 subunit suggesting the immunodominance of S1-RBD over S1-NTD region. This indicates that future antibody selection efforts to develop blocking antibodies directed towards non-RBD regions should be conducted with recombinant constructs lacking RBD. EC_50_ binding ELISA revealed 47 low- and sub-nanomolar binders to the S1-RBD domain. In the following ELISA and cell-based neutralization assays, 10 of the tested VNAR-hFc antibodies were able to disrupt engagement of S1 and S1-RBD to the ACE2 receptor. This inhibitory effect was confirmed for all 10 clones tested in a semi-quantitative assay in which Vero CCL81 cells were exposed to live SARS-CoV-2 virus of the Wuhan strain. Results of this assay showed good correlation with the preceding *in vitro* biochemical and cell-based assays. The live virus neutralization data was produced using VNAR-hFc antibodies at 5, 10, and 50 μg/ml (65, 130, and 650 nM) concentrations with comparable blocking activities. Future experiments could further examine the neutralizing potency of VNARs using serially diluted VNAR-hFc antibodies in the live virus assay.

Low resolution epitope binning was performed by competition ELISA and revealed two non-overlapping epitope bins in the S1-RBD domain recognized by VNAR-hFc antibodies. Of the 10 antibodies tested, 9 competed against each other for the overlapping epitopes and only one (3ID10_99) occupied a separate epitope bin. While the assay helped to identify the presence of at least two epitopes targeted by VNAR-hFc antibodies, the fact that the antibodies from a shared bin showed unique specificity profile for S1-RBD and mutants and unique blocking activity *in vitro* strongly suggests the tested VNAR-hFc antibodies occupied multiple epitopes. The observed competition in the ELISA binning assay was most likely affected by steric hindrance and would require a follow up assessment with high resolution approaches, such as alanine scanning mutagenesis.

A recent development in COVID-19 pandemic is the emergence of distinct phylogenetic virus clusters in geographically distant regions. RNA viruses, including SARS-CoV-2 are prone to mutations in the viral genome during replication. Some mutations are advantageous, allowing the virus to transmit more efficiently or escape the neutralizing effects of antibodies and other components of the host immune system. SARS-CoV-2 lineages in circulation were suggested to accumulate 1-2 nucleotide mutations per month (35). Studies indicated the accumulation of high mutation rates, including deletions and non-synonymous nucleotide changes, particularly in chronically infected COVID-19 patients with a deficient or suppressed immune system (36, 37). Presently most of the newly reported cases are attributed to 3 novel variants of the virus, lineage B.1.1.7/501Y.V1 (Kent), lineage B.1.351/501Y.V2 (South African), and lineage P.1 (Brazilian). The three clusters share some of the mutations linked with viral escape, including N501Y (B.1.1.7, B.1.351 and P.1), E484K (B.1.351 and P.1) and K417N/T (B.1.351 and P.1). Neutralizing antibodies targeting S1-RBD are broadly grouped into 4 classes (38). Antibodies in class 1 mainly form contacts around residues K417 and N501 and are therefore liable to mutations at these sites. Class 2 antibodies interact with the S1-RBD residue E484, as a result the E484K mutation confers resistance to this class of neutralizing antibodies (31, 39–42).

Given that the VNAR-hFc antibodies here were selected against S1 and S1-RBD derived from the ancestral SARS-CoV-2 (Wuhan strain), we wanted to understand how they would perform against the newly detected variants of the virus. The N501 residue is part of the binding loop of S1-RBD at the interface with human ACE2, forming a hydrogen bond with Y41 and stabilizing K353 on human ACE2 (43–46). The N501Y mutation was shown by deep mutation scanning to enhance binding affinity to ACE2 and to increase infectivity and virulence in a mouse model (47, 48). Our results showed that the N501Y mutation in S1-RBD had no effect on binding and blocking ability of the majority of tested VNAR-hFc antibodies. Only 3ID10_16 showed a 90-fold increase in EC_50_ binding affinity, and clones 3ID10_40 or 3ID10_96 each showed about 20-fold reduction or increase in IC_50_ blocking values, respectively.

The E484 residue of S1-RBD that directly interacts with K31 on ACE2 modestly enhances binding affinity to ACE2 (48). The E484K mutation was reported to abolish one of the salt bridges otherwise formed between ACE2 and S1-RBD (49). This mutation has been increasingly associated with reduced efficacy of vaccines and therapeutic antibodies, particularly with the B.1.351 and P.1 variants (50–53). Of the 10 VNAR-hFc antibodies tested against S1-RBD E484K, 3 showed reduction in EC_50_ binding but without losing their IC_50_ blocking activity. Given high correlation between *in vitro* blocking assays using recombinant proteins and live virus neutralizing potential when performed with the Wuhan variant, it can be expected that some of the VNAR-hFc antibodies would retain their blocking activity when tested against live virus containing N501Y and E484K mutations.

In summary, our work presents a new and novel class of single-domain antibodies against SARS-CoV-2 that are highly specific, and able to neutralize the ancestral SARS-CoV-2. The VNAR-hFc antibodies were also able to block S1-RBD with N501Y or E484K mutations *in vitro*, thus are expected to retain neutralizing potential towards virus variants that carry those substitutions.

## Abbreviations

ACE2: angiotensin converting enzyme 2
CDR3: complementarity-determining region 3
HSA: human serum albumin
NTD: N-terminal domain
RBD: receptor binding domain
RT: room temperature
SEC: size-exclusion chromatography
VNAR: variable domain of new antigen receptors
S: spike protein
SARS-CoV-2: severe acute respiratory syndrome coronavirus 2
SDS-PAGE: sodium dodecyl sulfate polyacrylamide gel electrophoresis

## Acknowledgement

This work was supported in part by a grant from the Pennsylvania Department of Community and Economic Development under the COVID-19 Pennsylvania Discoveries Program: Responding to SARS-CoV-2 Through Innovation & Commercialization (CV19-VTT). We thank Professor Kurt Zatloukal (Medical University of Graz) for his continuous support. A COVID-19 dedicated fund was kindly provided by the rectorate and faculty of natural sciences of the Karl-Franzens-University of Graz, which is hereby greatly acknowledged.

## Author Contributions

AG characterized VNAR antibodies, analyzed data, and co-wrote the manuscript; CVP performed phage selections and screening; MD purified proteins; JR performed viral infection experiments; TG analyzed viral infectivity assay; AJK coordinated viral experiments and co-wrote the manuscript; FSW and JLR designed research and co-wrote the manuscript; PS designed and led the research and co-wrote the manuscript

